# Peptimetric: Quantifying and visualizing differences in peptidomic data

**DOI:** 10.1101/2021.05.18.444693

**Authors:** Erik Hartman, Simon Mahdavi, Sven Kjellström, Artur Schmidtchen

## Abstract

Finding new sustainable means of diagnosing and treating diseases is one of the most pressing issues of our time. In recent years, several endogenous peptides have been found to be both excellent biomarkers for many diseases and to possess important physiological roles which may be utilized in treatments. The detection of peptides has been facilitated by the rapid development of biological mass spectrometry and now the combination of fast and sensitive high resolution MS instruments and stable nano HP-LC equipment sequences thousands of peptides in one single experiment. In most research conducted with these advanced systems, proteolytically cleaved proteins are analyzed and the specific peptides are identified by software dedicated for protein quantification using different proteomics workflows. Analysis of endogenous peptides with peptidomics workflows also benefit from the novel sensitive and advanced instrumentation, however, the generated peptidomic data is vast and subsequently laborious to visualize and examine, creating a bottleneck in the analysis. Therefore, we have created Peptimetric, an application designed to allow researchers to investigate and discover differences between peptidomic samples. Peptimetric allows the user to dynamically and interactively investigate the proteins, peptides, and some general characteristics of multiple samples, and is available as a web application at https://peptimetric.herokuapp.com. To illustrate the utility of Peptimetric, we’ve applied it to a peptidomic dataset of 15 urine samples from diabetic patients and corresponding data from healthy subjects.

## I. Introduction

Although peptides have been studied since the beginning of the 20th century, the practice of analyzing the complete peptidome of large cohorts of samples has only recently been actualized by the advancement of biological mass spectrometers, tandem-mass spectrometry techniques (MS/MS) and computational algorithms [1]. These new techniques allow for large screening of endogenous bioactive molecules with great physiological implications which ultimately may give rise to effective and sustainable pharmaceuticals and diagnostic tools [2]. New and more advanced mass spectrometers generate progressively more data as they’re able to identify more peptides and perform searches faster [3]. Large sets of peptidomic data are difficult to comprehend and visualize without advanced and dedicated programs, and there is therefore a great need for tools which enable researchers with modest experience in programming and visualization skills to analyze their samples.

The nature of peptidomic data makes it highly compatible with computational algorithms and there are already many software programs which utilize this fact to e.g. map post translational identifications or screen for antimicrobial peptides [4]. One useful tool for visualization of multiple peptidomic samples is Peptigram [5], created by Manguy et al. Peptigram allows for easy peptide visualization using an internet based framework, and connects with various resources such as Pfam [6] and ProViz [7] to create a comprehensive map of the peptidome of multiple samples. However, as with every tool, there are some limitations to Peptigram, especially when it comes to pre-processing, quantitative comparisons between sample groups and interactive sample exploration.

The process of producing MS/MS data in data dependent mode is stochastic and results in variation between samples, and variation is also added by sampling techniques used prior to the liquid chromatography tandem-mass spectrometry (LC-MS/MS) analysis which leads to biases between samples. These biases are of extreme importance if one tries to find biomarkers, quantify patterns or statistically determine correlations. Therefore, peptidomic data regularly require pre-processing where normalization techniques and cutoffs are introduced, so that inter-sample biases are minimized and peptides of low abundance or certainty are removed from the dataset [8].

Furthermore, scientific hypothesis testing generally consists of groups of samples, where a difference or correlation of an observed phenomenon is quantified and its statistical significance determined. This is especially true of medical and biological trials, as many studies compare positive samples to a negative control group. Additionally, the nature of peptidomic data is highly dependent on the connection between precursor protein and the resulting peptides. To explore these relationships effectively a high level of interactiveness is required, where one can easily switch between investigating the identified specific endogenous peptides and the sequence coverage from the corresponding proteins.

In previous work, computation in combination with tools such as Peptigram was utilized to explore the presence of biomarkers and endogenous antimicrobial peptides in infected wounds, further showcasing the applicability and usefulness of algorithms on peptidomic data [9]. In this paper, we have continued to explore and develop algorithms allowing for massive screening and visualization of peptides in large cohorts of samples. Ultimately, we’ve created Peptimetric, an open-source web based application for highly interactive group comparisons and sample exploration of peptidomic data, available at https://peptimetric.herokuapp.com/. Peptimetric allows the user to pre-process their data using different types of normalization and cutoffs to remove biases which are introduced in sample preparation and subsequent LC-MS/MS analysis. Thereafter it provides a dynamic graphical interface where the user may investigate their sample groups on a protein and peptidomic level to search for peptides of interest. Furthermore, the generated figures and tables are readily available to download for publications or further exploration. Peptimetric is freely available as an open GitHub repository (https://github.com/ErikHartman/peptimetric) under an MIT license.

## II. Method

### A. Implementation

Peptimetric is implemented as a web application making it accessible through a web browser without the need for package or software-installations at (https://peptimetric.herokuapp.com/). The application is hosted by the cloud service Heroku (https://www.heroku.com/). The front end of the application is developed in Python 3.8.8 using the Dash library (version 1.2.0) (https://plotly.com/dash/) in combination with Bootstrap components. Peptimetric is therefore compatible with the following web browsers: Chrome, Opera, Microsoft Edge and Firefox and Safari. Note that initial loading of the application may be lengthy due to the nature of the Heroku platform and our subscription plan (https://devcenter.heroku.com/articles/free-dyno-hours#dyno-sleeping). The complete repository, including the data files and requirements, are available at GitHub (https://github.com/ErikHartman/Peptimetric) and is freely available under an MIT license [10]. The repository may be cloned to run Peptimetric locally.

### B. Input files

Peptimetric is used to visualize and dynamically explore the differences between the peptidomes of groups of samples. Therefore, at least one data-file, in a .csv (comma separated values) format, per group is required to use the application. However, for statistical analysis, a minimum of three files per group is required. The files need to include four columns for the peptide sequence, intensity (or other abundance metric), spectral count (or other abundance metric) and a protein precursor id (UniProt id). To accommodate for different raw data processing softwares, such as PEAKS, the names in table 1 (including capitalization variations) are accepted for the columns:

**Table I.** Allowed column names for the input files.

|  |  |
| --- | --- |
| Peptide sequence | Peptide, Sequence |
| Precursor protein ID | Accession, Protein, UniProt ID |
| Intensity | Area, Intensity |
| Spectral count | SpC, Spectral count, #Feature |

All input files are stored on the Heroku server during the session. A local copy of proteomes fetched from UniProt TrEMBL in April of 2021 is used to get the FASTA sequences and UniProt mnemonic protein identifier (e.g. HBA_HUMAN) for the submitted precursor proteins [11]. The local database consists of a subset of proteomes which were chosen due to their prevalence in biological and peptidomic studies (see table II). Additional proteomes are easily added, and we urge researchers to contact us if they wish to analyze samples from other species.

**Table II.**
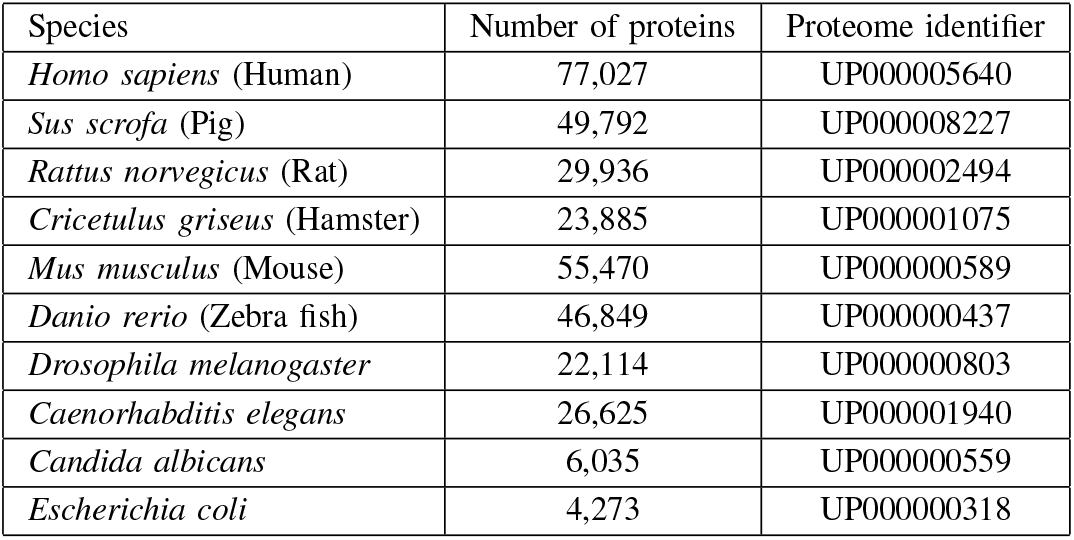
The species currently available in the Peptimetric database, alongside the number of proteins and UniProt proteome identifier (UPID).

### C. Data processing

The first step of data manipulation performed by Peptimetrics is to take the 10th logarithm (log10) of all intensity values. This is commonly performed to diminish the exponential nature of MS intensities and to later provide appropriate quantification for proteins and peptides as Gaussian (normal) distribution is assumed [12]. If one has already taken the logarithm of their values, or wishes to leave the intensity values unaltered, a checkbox in the normalization popup is to be checked before uploading the files.

Biases are often introduced during sample preparation LC-MS/MS analysis [8], [13]. Some of these biases may be dealt with when processing the raw files, however, they may go undetected or one may choose to leave the samples unaltered to perform post-processing normalization. Additionally, peptidomic data may include peptides and proteins with low abundance or certainty which may not be appropriate to include in peptidomic analysis. Therefore, Peptimetric includes ways to normalize the input data, as well as to discard sets of the data based on cutoffs.

Normalization of the data from MS or MS/MS can be performed in multiple ways, however, Peptimetric accommodates for two ways of normalization: either by normalizing on the samples’ global values [14], [15] or by normalizing on a housekeeping protein [16]. When normalizing on the global values the intensity/spectral count for each peptide is multiplied by the mean sum of intensities/spectral count divided by the sum of the intensity/spectral count respectively in the sample, see equation (1).

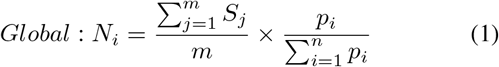

where:

N normalized value

S total sample value

m number of samples

p peptide value

n number of peptides in sample

Normalization by a housekeeping protein is performed similarly to the global normalization, however, instead of using the total sum based on the complete sample, a subsection containing peptides from the selected precursor protein is used for normalization, see equation (2).

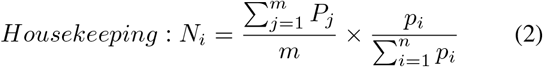

where:

N normalized value

P total protein value

m number of samples

p peptide value

n number of peptides in protein

If the housekeeping protein is not present in a given sample, the intensity and spectral count values in the sample are left unaltered.

Cutoffs are used to remove proteins and/or peptides below given thresholds and are applied to peptides and proteins separately. Peptides may be discarded based on intensity, spectral count and for being retention time (RT) and/or collision cross section (CCS) outliers. Outliers are defined to be situated three standard deviations from the mean value. Protein cutoffs may thereafter be applied to remove proteins with cumulative intensities or spectral counts below a given threshold. Additionally, proteins containing several peptides below a given threshold may be removed. Note that protein cutoffs are applied after peptide cutoffs. Therefore, proteins which contain many peptides of low abundance will be removed if the peptide cutoffs are applied appropriately.

### D. Experimental data

To illustrate the utility and typical usage of Peptimetric, a dataset generated from a study by Van et al, 2020, describing peptidomic analysis of urine, was fetched from an online repository [17]. The study profiles the urinary peptides from 15 patients with type-1-diabetes (D) and uses corresponding data from non-diabetic (ND) subjects for comparison. This study is well suited for analysis using Peptimetric as it contains groups of samples and uses a discovery peptidomics approach. Additionally, the study collected data from a relatively large population and conducted thorough qualitative and quantitative analyses. Details about study population, the sample generation and preparation are described further in Van et al [17].

The raw files were retrieved from ProteomeXchange Consortium via the PRIDE partner repository with the dataset identifier PXD012210 (http://proteomecentral.proteomexchange.org/cgi/GetDataset?ID=PXD012210) They were then searched using PEAKS Xpro with similar settings using a maximum of 2 modifications per peptide, allowing the same modifications (methionine and proline oxidation, N-terminal acetylation). After the database search the identified peptides were filtered with a cut off using 1% FDR and a minimum of two peptides per protein. In total, 6559 and 9024 unique peptides were identified in the samples from non-diabetics and type-1-diabetics respectively. The files are uploaded to the server, and the data may be investigated by any user by clicking “Load Example Files”.

The complete workflow is described in figure 1 below.

**Figure 1.**
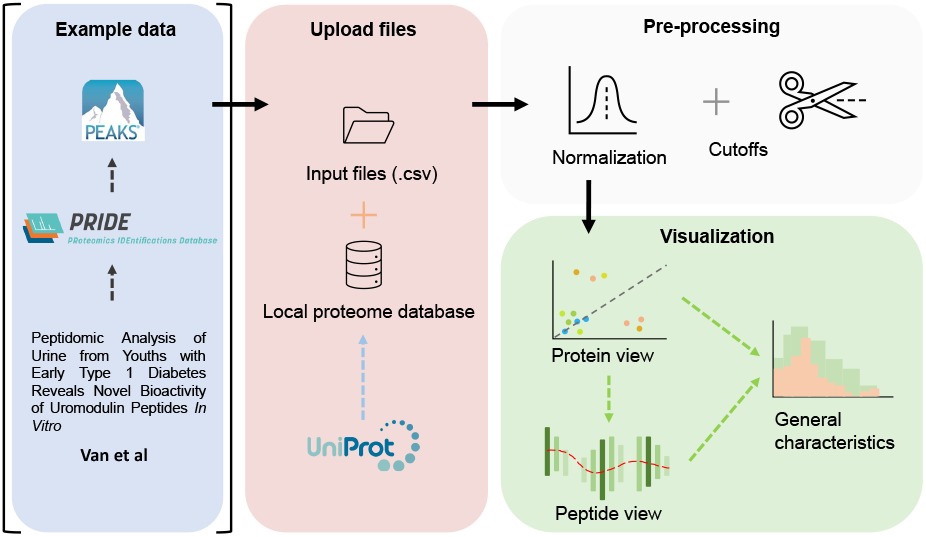
An overview of the workflow performed in this study. The raw files, deposited by Van et al. [17], were fetched from ProteomeXchange Consortium via the PRIDE partner repository, and searched against a human proteome database using PEAKS. In Peptimetric, the input files are matched against a local proteome database which was fetched from UniProt. Normalization and cutoffs are applied to the dataset. The data is visualized in a protein view, which in turn may be used to generate a peptide view. An overview of some general characteristics may also be generated for either the complete proteome or the selected protein.

## III. Result

After submitting input files into each group, the files are concatenated and matched against the local database to retrieve the FASTA sequences and UniProt mnemonic identifiers. If a submitted UniProt id is not found in the database, the peptides belonging to the precursor protein will be discarded from the dataset.

### A. Protein view

After uploading the files and applying normalization and cutoffs, it is possible to generate a protein graph which showcases all the precursor proteins present in the samples in the format of a scatter plot, see figure 2A. The protein graph allows the user to visualize the abundance of the proteins based on one of the following metrics: sum of intensities, mean of intensities, sum of the spectral counts, and mean of the spectral counts. The means are calculated by taking the mean of the abundance metric of all the peptides in each protein, whereas the sum is calculated by taking the sum of the abundance metric of all the peptides in each protein, for each individual sample. Thereafter, the resulting values are averaged across the samples. This results in a value for each group, which is plotted as X- and Y-coordinates in the scatter plot. Alongside with the abundance metric chosen by the user the protein graph also provides another two dimensions of visualizing differences between the sample groups, namely: the number of peptides derived from each precursor protein, which correlates to the dot size, and the absolute value of the difference between the chosen abundance metric between the groups, which is proportional to the color. Since the protein abundance for each group is plotted along the X- and Y-axis, the diagonal represents the point of equal abundance between the groups. Furthermore, the standard deviation of the abundance metric is off by default but can be displayed in the protein graph.

**Figure 2.**
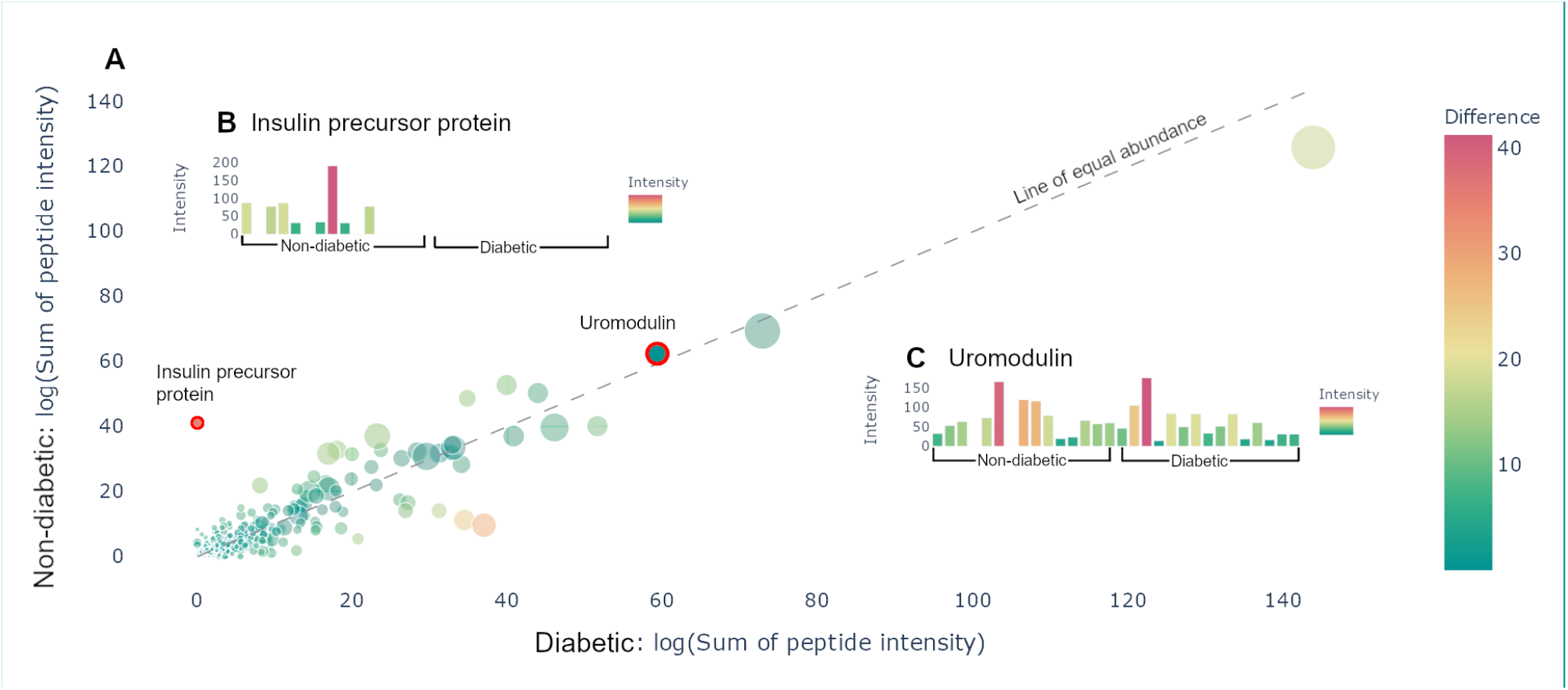
Protein view. **A** Scatter plot containing all the proteins present in the samples is presented. The size of the dots corresponds to the number of peptides derived from the precursor protein. The color corresponds to the distance from the line of equal abundance, i.e., the difference in abundance metric between the two groups. **B** Hovering over the insulin precursor protein results in the presented sample graph. Peptides are only present in non-diabetics. **C** Hovering over uromodulin generates the presented sample graphs. Uromodulin is varyingly present in both groups.

The protein graph may be manipulated in multiple ways. All graphs created in Peptimetric are generated with Plotly’s graphing tools, which provides the user with a built in modebar containing tools for e.g., downloading the plots (https://plotly.com/python/configuration-options/). Hovering over a protein in the protein graph displays the metrics for that specific protein. Simultaneously, hovering over a protein will produce a sample graph (see figure 2B,C), containing the abundance metric for each individual sample for the protein.

All proteins present in the samples are summarized in a protein table (see table III. The protein table contains the UniProt mnemonic protein identifier, UniProt id, number of peptides, abundance metric value, the standard deviation and the p-value (although the p-value is only shown if the given protein is present in three or more samples in each group) for the given abundance metric between the two groups. The table may be filtered with regular expression syntax, e.g.: >, ≤, ≥, = (for more see https://dash.plotly.com/datatable/filtering).

**Table III.**
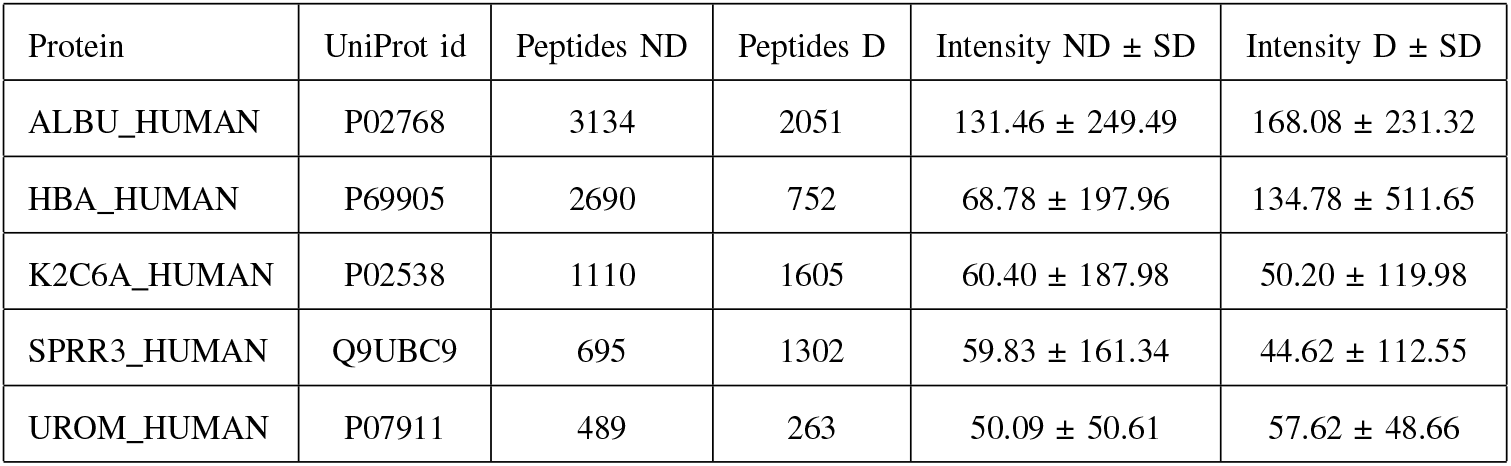
Protein table. The table contains the top 5 most abundant proteins in both groups.

| Protein | UniProt id | Peptides ND | Peptides D | Intensity ND ± SD | Intensity D ± SD |
| --- | --- | --- | --- | --- | --- |
| ALBU_HUMAN | P02768 | 3134 | 2051 | 131.46 ± 249.49 | 168.08 ± 231.32 |
| HBA_HUMAN | P69905 | 2690 | 752 | 68.78 ± 197.96 | 134.78 ± 511.65 |
| K2C6A_HUMAN | P02538 | 1110 | 1605 | 60.40 ± 187.98 | 50.20 ± 119.98 |
| SPRR3_HUMAN | Q9UBC9 | 695 | 1302 | 59.83 ± 161.34 | 44.62 ± 112.55 |
| UROM_HUMAN | P07911 | 489 | 263 | 50.09 ± 50.61 | 57.62 ± 48.66 |

The user is also able to search for all proteins present in the samples, that have not been removed by any potential cutoffs. Searching for, clicking on, or selecting a protein in the table highlights it in the protein graph, making it possible to create a peptide graph for the highlighted protein.

### B. Peptide view

Once a protein is selected in the protein view, a peptide view may be generated to view the peptides of the selected protein. Doing so results in a peptide graph alongside a peptide table. The peptide graph follows a visualization convention where the peptides are mapped onto the complete protein sequence as barcharts. To allow for easy visualization and quantification of peptide abundance and coverage, the color of the bars is proportional to the number of overlapping peptides at each amino acid position, whereas the height of the bars is proportional to the used abundance metric, see figure 3. Similarly to the protein view, spectral count and intensity are available abundance metrics, and the user may easily switch between the two.

**Figure 3.**
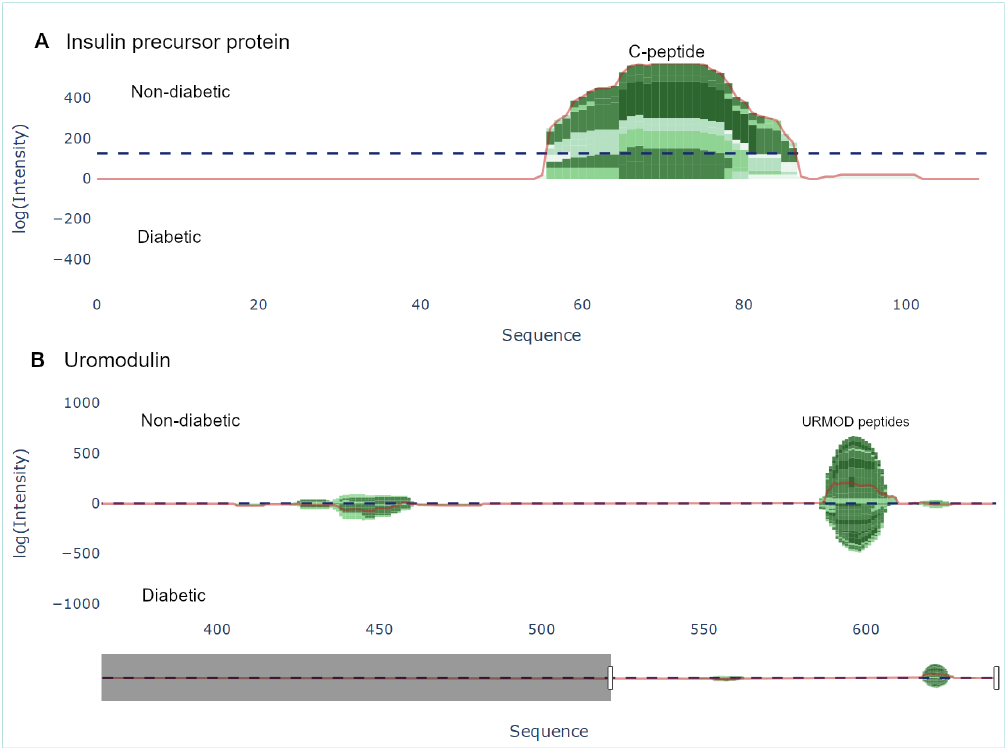
Peptide view of insulin precursor protein (INS_HUMAN) and uromodulin (UROM_HUMAN). **A** The C-peptide (57-87) from the insulin precursor protein is present in group 1 (non-diabetic). **B** There are two distinct regions present in uromodulin. The second region (430-460) contains the peptides UMOD1-UMOD7 and was documented by Van et al. [17].

There are two options for graphically representing the peptidome of the input files: either as individual samples, or by averaging each group. If viewing each individual sample, the samples are stacked, whereas when viewing the group average, the intensity at each position is averaged. To guide the user to interesting regions of difference, a line corresponding to the difference of the height at each position is plotted. Additionally, a horizontal line representing the weight, i.e. the sum of the chosen abundance metric. All the components in the graph are displayed as individual traces, and may be hidden/shown.

A table showing the peptides in the samples is situated adjacent to the peptide graph. The table shows the peptide amino acid sequence, the start and end position of the sequence, as well as the group mean, standard deviation and p-value of the chosen abundance metric (see table IV). The table is filterable and sortable in a similar manner as the protein table. Each sequence in the table is selectable and by doing so highlights the peptide region in the peptide graph.

**Table IV.** Peptide table for uromodulin (UROM_HUMAN). The table contains the 5 most abundant peptides in both groups.

| Peptide | Start | End | Intensity ND ± SD | Intensity D ± SD |
| --- | --- | --- | --- | --- |
| SGSVIDQSRVLNLGPITRKG VQ | 588 | 610 | 4.68 ± 14.16 | 0.59 ± 2.30 |
| VGGTGMFTVRM | 449 | 460 | 2.39 ± 5.31 | 2.11 ± 5.70 |
| SGSVIDQSRVLNLGPITRK | 588 | 607 | 2.24 ± 6.00 | 0.18 ± 0.71 |
| VGGTGMFTVRMA | 449 | 461 | 1.95 ± 5.17 | 2.18 ± 5.09 |
| LQPMVSALNIR | 438 | 449 | 1.62 ± 4.42 | 2.14 ± 5.70 |

### C. General characteristics

The dynamics of the peptidome are largely due to enzymatic activity, resulting in variations in peptide length and amino acid profile of the peptides N- and C-terminals. Therefore, Peptimetric includes three relevant general characteristics, which may be applied to either the peptides in the entire proteome or in the chosen protein. These include: a histogram over peptide length, a bar chart showing peptide overlap between the groups, and pie charts displaying the amino acid profile, see figure 4.

**Figure 4.**
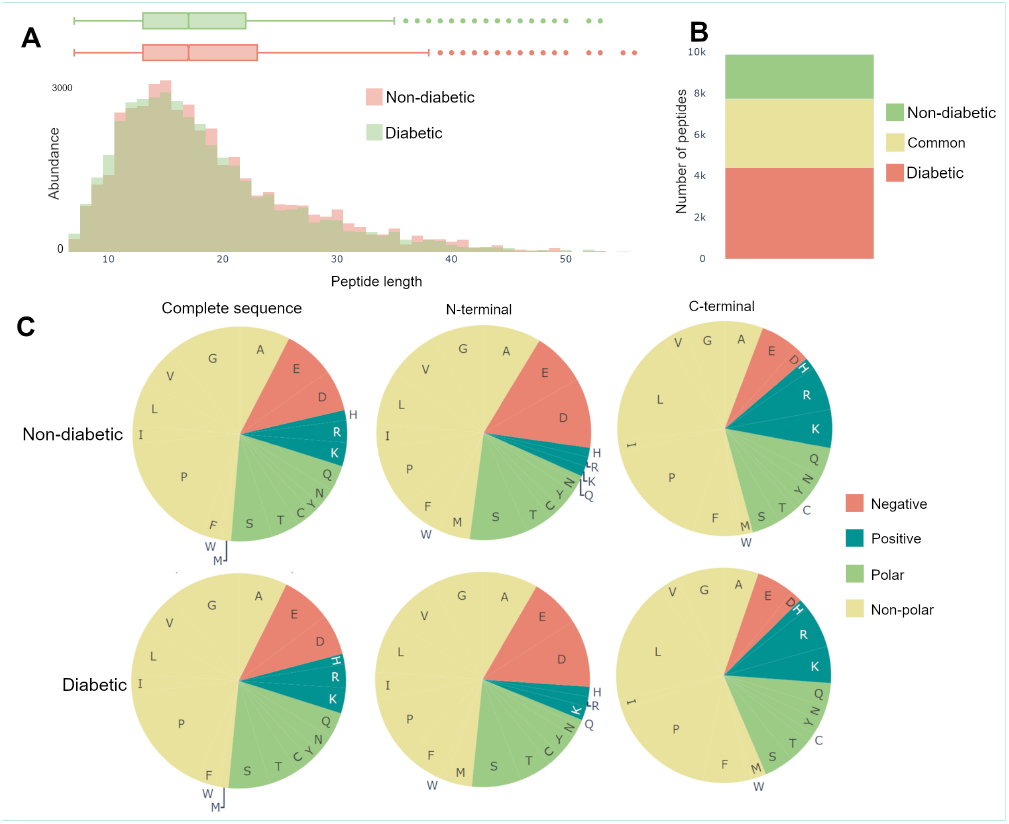
General characteristics of the complete peptidomes. **A**. The length distribution of peptides, weighted by the abundance metric. **B**. Venn bars, showing the overlap of the peptidomes. **C**. Amino acid profile, showing the amino acid prevalence for the complete sequence, as well as the first and last amino acid.

The length distribution is created by weighting each peptide with the value of the specific abundance metric, and thereafter counting the number of peptides with each length. Therefore, if spectral count is chosen as the abundance metric, a peptide with the spectral count value of 10 will contribute twice as much to the length distribution as a peptide with a spectral count value of 5.

The amino acid profiles contain pie charts showing the distribution of amino acids for the complete amino acid sequence, the first amino and the last amino acid, as can be seen in figure 4. The abundance of an amino acid is calculated similarly to the length distribution, as described above. The color scheme for the pie charts is created based on the polarity and acidity of the amino acids [18].

### D. Exploring the dataset

Investigating the data yielded an overview of the most abundant proteins in the samples. After normalizing on global values and using the sum of intensities as the abundance metric, the most abundant proteins were albumin, hemoglobin alpha, uromodulin, small proline-rich protein 3 (SPRR3) and keratin type II cytoskeletal 6A, which is in line with the results from Van et al [17]. Additionally, apolipoprotein A1 and alpha-1-antitrypsin (A1AT) were noticeably more abundant in the diabetic samples, whereas insulin precursor protein is only present in non-diabetic samples (see figure 2A. It is worth to note that the samples vary a lot, and no difference is statistically significant on the protein level. Similar conclusions were drawn with all different abundance metrics.

Van et al. [17] describes 7 C-terminal peptides (UMOD1-UMOD7, most significantly SGSVIDQSRVLNLGPITRK, 588-606) from uromodulin as biomarkers for type-1-diabetes. When investigating the dataset, many peptides derived from this region were indeed identified (alongside the region 430-460), however, only modest differential expressions between the two groups were present, see figure 3. As mentioned above insulin precursor protein was not found in the samples taken from the patients with type-1 diabetes. Investigating insulin precursor protein in the peptide view showcases that all peptides are derived from the region 57-87, which corresponds to the well documented C-peptide [19]. Although there was only modest overlap between the peptidomes of the groups (30%), there were no apparent differences regarding peptide length or amino acid profile between the groups (see figure 4).

### E. Limitations

Peptimetric uses a local database fetched from UniProt to process the input data to reduce loading times. This results in the disadvantage of limiting the number of species accessible for analysis as well as not having the latest version of the UniProt database. The size of the local database itself is limited by the Heroku slug storage size, which is restricted to 500 MB for the subscription plan (https://devcenter.heroku.com/articles/slug-compiler#slug-size).

Herokus servers only allow for a limited computation time to each process (30 seconds), and processes which surpass the limit are aborted. This may occur if the user tries to input several large files with a slow internet connection. If this occurs, concatenating some of the files manually is recommended, since this will reduce processing time.

Despite these limitations, Peptimetric serves as an interactive platform for fast and user-friendly exploration of MS/MS data for detailed investigations and visualization of complex data sets.

## IV. Conclusion

Peptimetric allows researchers to effectively and dynamically process, investigate and explore their peptidomic data. These traits are highly sought after in a landscape where the throughput of data is ever growing and where computational algorithms play an increasingly important role. By implementing an interface where the user easily gets an overview of both the proteome and peptidome of a sample, we have created a tool with high applicability to various peptidomic projects. To illustrate the utility of Peptimetric, we applied it to a dataset generated by Van et al, [17] where the urinary peptidomes of type-1-diabetics were studied.

We envision Peptimetric being used in the early stages of peptidomic data analysis, where overview, exploration and quantification play important roles in the identification of precursor proteins and peptides of interest. Thereafter, one may export the data retrieved from Peptimetric for further analysis of e.g., post translational modifications, enzyme cut sites and screenings for bioactive peptides. This methodology applies to a variety of possible studies as peptidomics has relevance in basic research and clinical studies.

## V. Acknowledgements

We would like to thank Arvid Larsson, who supported and advised us through the entirety of this project and did not demand nor want anything in return.

## VI. Author Contribution

EH and SM wrote the complete manuscript, which was revised by SK and AS. EH and SM wrote the code. EH, SM, SK and AS developed the ideas behind Peptimetric.

